# Defects of myelination are common pathophysiology in syndromic and idiopathic autism spectrum disorder

**DOI:** 10.1101/128124

**Authors:** BaDoi N. Phan, Stephanie Cerceo Page, Morganne N. Campbell, Joseph F. Bohlen, Courtney L. Thaxton, Jeremy M. Simon, Emily E. Burke, Joo Heon Shin, Andrew J. Kennedy, J. David Sweatt, Benjamin D. Philpot, Andrew E. Jaffe, Brady J. Maher

## Abstract

Autism Spectrum Disorder (ASD) is genetically heterogeneous in nature with convergent symptomatology, suggesting dysregulation of common molecular pathways. We analyzed transcriptional changes in the brains of five independent mouse models of Pitt-Hopkins Syndrome (PTHS), a syndromic ASD caused by autosomal dominant mutation in *TCF4,* and identified considerable overlap in differentially expressed genes (DEGs). Gene and cell-type enrichment analyses of these DEGs identified oligodendrocyte dysregulation that was subsequently validated by decreased protein levels. We further showed significant enrichment of myelination genes was prevalent in two additional mouse models of ASD (*Pten^m3m4/m3m4^, Mecp2^KO^*). Moreover, we integrated syndromic ASD mouse model DEGs with ASD risk-gene sets (SFARI) and human idiopathic ASD postmortem brain RNA-seq and found significant enrichment of overlapping DEGs and common biological pathways associated with myelination and oligodendrocyte differentiation. These results from seven independent mouse models are validated in human brain, implicating disruptions in myelination is a common ASD pathophysiology.

## Main Text

Autism spectrum disorder (ASD) affects approximately 1:68 individuals and has incalculable burdens on affected individuals, their families, and health care systems. While the genetic contributions to idiopathic ASD are heterogeneous and largely unknown, the causal mutations for syndromic forms of ASD – including truncations and copy number variants – provide a genetic toehold with which to gain mechanistic insights(*1*-*3*). Models of these syndromic disorders have been used to better characterize the molecular and physiological processes disrupted by these mutations(*4*). Two fundamental questions remain – how biologically similar are the mouse models of syndromic forms of ASD, and how relevant are these mouse models to their human analogs? To address these questions, we performed integrative transcriptomic analyses of seven independent mouse models of three syndromic forms of ASD generated across five laboratories, and assessed dysregulated genes and their pathways in human postmortem brain from patients with ASD and unaffected controls. These cross-species analyses converged on shared disruptions in myelination and axon development across both syndromic and idiopathic ASD, highlighting both the face validity of mouse models for these disorders and identifying novel convergent molecular phenotypes amendable to rescue with therapeutics.

We first assessed molecular convergence across five different mouse models of Pitt-Hopkins syndrome (PTHS), a rare form of ASD that results from diverse mutations in the transcription factor 4 (*TCF4)* gene, ranging from haploinsufficiency to dominant-negative. This syndromic disorder is characterized by developmental delay, failure to acquire language, motor learning deficits, and gastrointestinal abnormalities(*5*-*7*). We generated RNA-seq data from prefrontal cortex (PFC) of a PTHS mouse line that shows heterozygous expression of a truncated TCF4 protein (*Tcf4*^+/tr^)(*8*) (see Methods). The *Tcf4*^+/tr^ mouse showed significant blunted expression of full-length *Tcf4* transcript (p=0.02, Fig. S1A, B) and protein (p=0.0009, Fig. S1C, D) in the brain between embryonic day 16 and postnatal day 4 (E16, P4) and smaller difference in expression in adulthood (p>0.05, Fig. S1A, B, D). A similar expression pattern was observed across the human lifespan(*9*), suggesting there may be a critical period for the genesis of PTHS that is concordant with early cortical development, a critical causal period identified for other syndromic and idiopathic forms of human ASD(*10*, *11*). In addition to the *Tcf4*^+/tr^ PFC RNA-seq and published of *Tcf4*^+/tr^ hippocampal CA1(*12*) dataset, we further created and processed RNA-seq data from four additional mouse lines of heterozygous *Tcf4* mutations or deletions: *Tcf4*^+/Δ574-579^, *Tcf4*^+/R579W^, *Actin-Cre*::*Tcf4*^+/floxed^ and *Nestin*-*Cre*::*Tcf4*^+/floxed^ (see Methods). Given that *Tcf4* is developmentally regulated and plays a role in gene regulation, we assessed the effects of heterozygous *Tcf4* mutations (*Tcf4*^+/mut^) on the mouse transcriptome from the prefrontal cortex, CA1 region of hippocampus, and hemi-brain at P1 and in adulthood (see Fig. 1A for details).

**Fig. 1:**
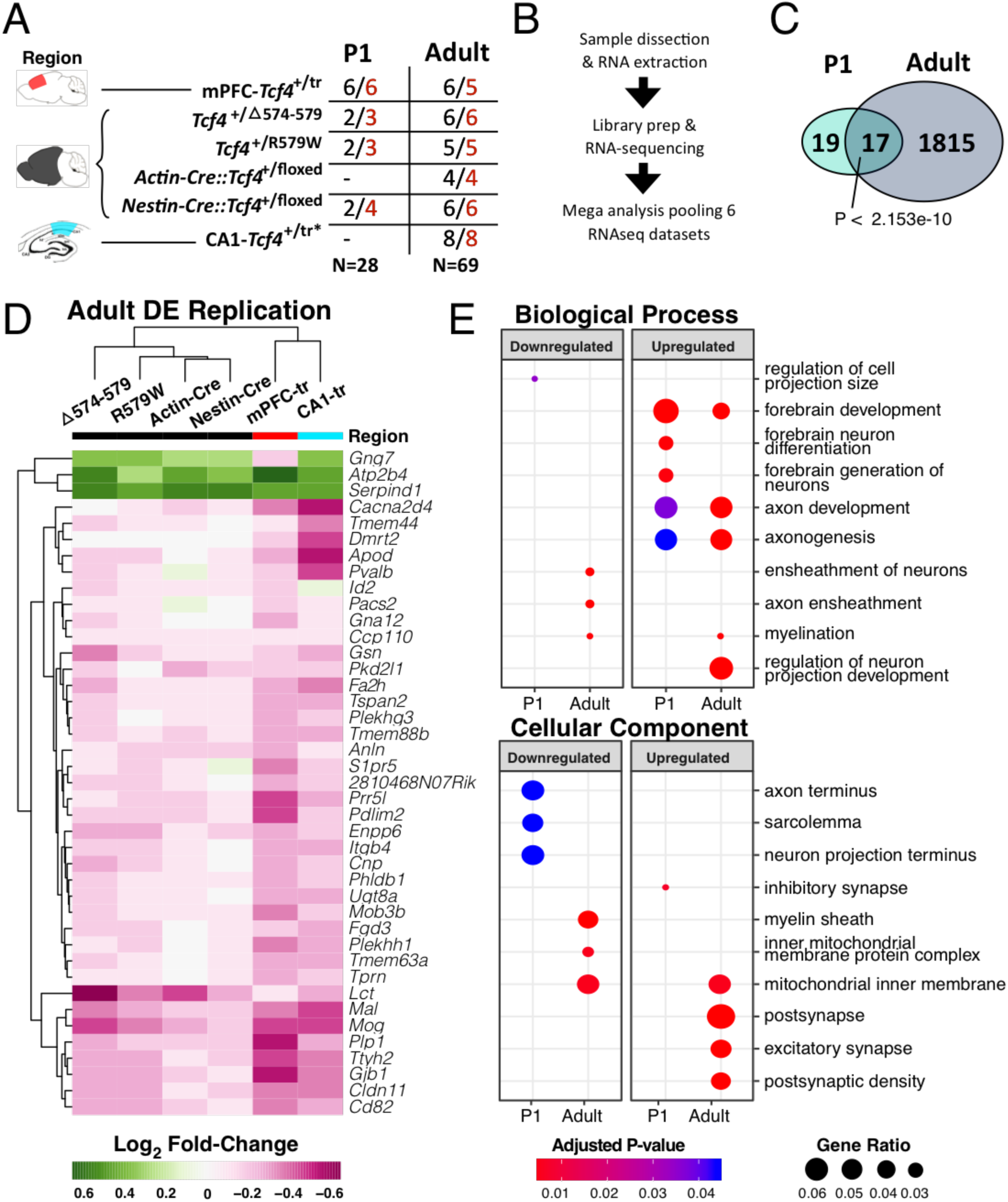
RNA-seq of multiple *Tcf4* mutations reveal age-specific differential gene expression. **(A)** Summary table of the 5 mouse lines of *Tcf4* mutation sequenced in this analysis. Samples come from 3 regions, medial prefrontal cortex, hemibrain, and hippocampal CA1 (colored red, black, and teal, respectively)(*12*). Two age groups, P0-2 (P1) and >P42 (adult), were assessed in this study. N’s of wild-type and *PTHS* mice are colored black and red, respectively. **(B)** General sample-to-analysis RNA-seq pipeline. **(C)** Venn diagram of differentially expressed genes (DEGs) in P1 and adult mice by *Tcf4*^+/mut^ genotype across all mouse lines and tissue regions (FDR<0.05). There are 36 DEGs in P1 group, and 1832 DEGs in adult group. A significant group of 17 genes are differentially expressed in both P1 and adult age groups (p<2e-10). **(D)** Log_2_ fold-change heatmap of DEGs from the mega analysis shows high concordance of differential expression across mouse lines (replication p<0.05). **(E)** Dot plot of gene ontology (GO) enrichment analysis of marginal DEGs split by up-or down-regulated genes to determine functional pathways affected in *PTHS* mice brain (P_adj_ < 0.05). Gene ratio dot size represent % of genes for each GO term differentially expressed.

Differential expression analysis of each *Tcf4*^+/mut^ mouse model by age showed more differentially expressed genes (DEGs) in the adult brain than in the P1 brain, with overall high concordance and replication rates across these varied models and tissue sources (Fig. 1D, S2, Table S1). Combined analysis across the RNA-seq data from these multiple models of PTHS (“mouse mega-analysis”) revealed widespread transcriptional dysregulation in the adult mouse brain in *Tcf4*^+/mut^ versus wild-type (WT) (Fig. 1B, Table S2). Gene ontology (GO) analysis of the DEGs showed age-specific processes involved in *Tcf4*^+/mut^ (Fig. 1E, see Table S3 for full GO analysis) – genes upregulated in adult brains were enriched for processes including forebrain development, neuron projection, axon development, excitatory and inhibitory synapse, postsynaptic density, and transcription cofactor activity while those genes downregulated enriched for neuron ensheathment, myelination, and peptide binding. Cell type-specific analysis (CSEA) using expression levels from microarrays(*13*) (p=2E-26, Fig. S3) and RNA-seq of purified mouse cell types(*14*) further implicated oligodendrocytes as the candidate cell type implicated in *Tcf4*^+/mut^ (Fisher p_adj_<0.05, Fig. 2A). For example, adult mouse lines across all *Tcf4*^+/mut^ models showed oligodendrocyte enrichment at all developmental stages with the most significant enrichment found in myelinating oligodendrocytes, where more than 55% of cell type-specific genes were differentially expressed in our mega-analysis. We further applied cellular deconvolution to obtain the estimated proportion of each cell type and identified a significant increase in the proportions of neurons (p=0.035) and astrocytes (p=0.007), and significant decrease in the proportions of myelinating oligodendrocytes (and p=0.002) in adult *Tcf4*^+/mut^ mice. From previous purified cell type-expression data(*14*), we see that *Tcf4* is moderately expressed in oligodendrocyte precursor cells and newly formed oligodendrocytes, but lowly expressed in myelinating oligodendrocytes, suggesting decreased *Tcf4* expression in immature cell types may obstruct oligodendrocyte maturation (Fig. 2B). To validate that these mature oligodendrocyte-specific RNA-seq signatures reflect functional deficits in our PTHS mouse models, we measured the expression of key oligodendrocyte proteins (CNP, MOG, and NG2) from *Tcf4*^+*/tr*^ and WT adult brain lysates (Fig. 3D). Relative protein levels of myelinating oligodendrocyte markers, CNP and MOG, were significantly decreased in *Tcf4*^+*/tr*^ mice (Fig. 3E, N=12 each, p_MOG_=0.0008, p_CNp=_0.0436) and their fold-changes in mutant vs. WT mice correlated with fold-changes from the RNA-seq differential expression analysis (Fig. 3F). However, the relative protein level and RNA-seq expression of NG2, the marker for oligodendrocyte precursor cells, was not different between genotypes (N=12, p=0.815). Together, these data suggest that TCF4 is critical to the proper maturation of oligodendrocytes and the process of myelination but not in the development of oligodendrocyte precursor cells.

**Fig. 2:**
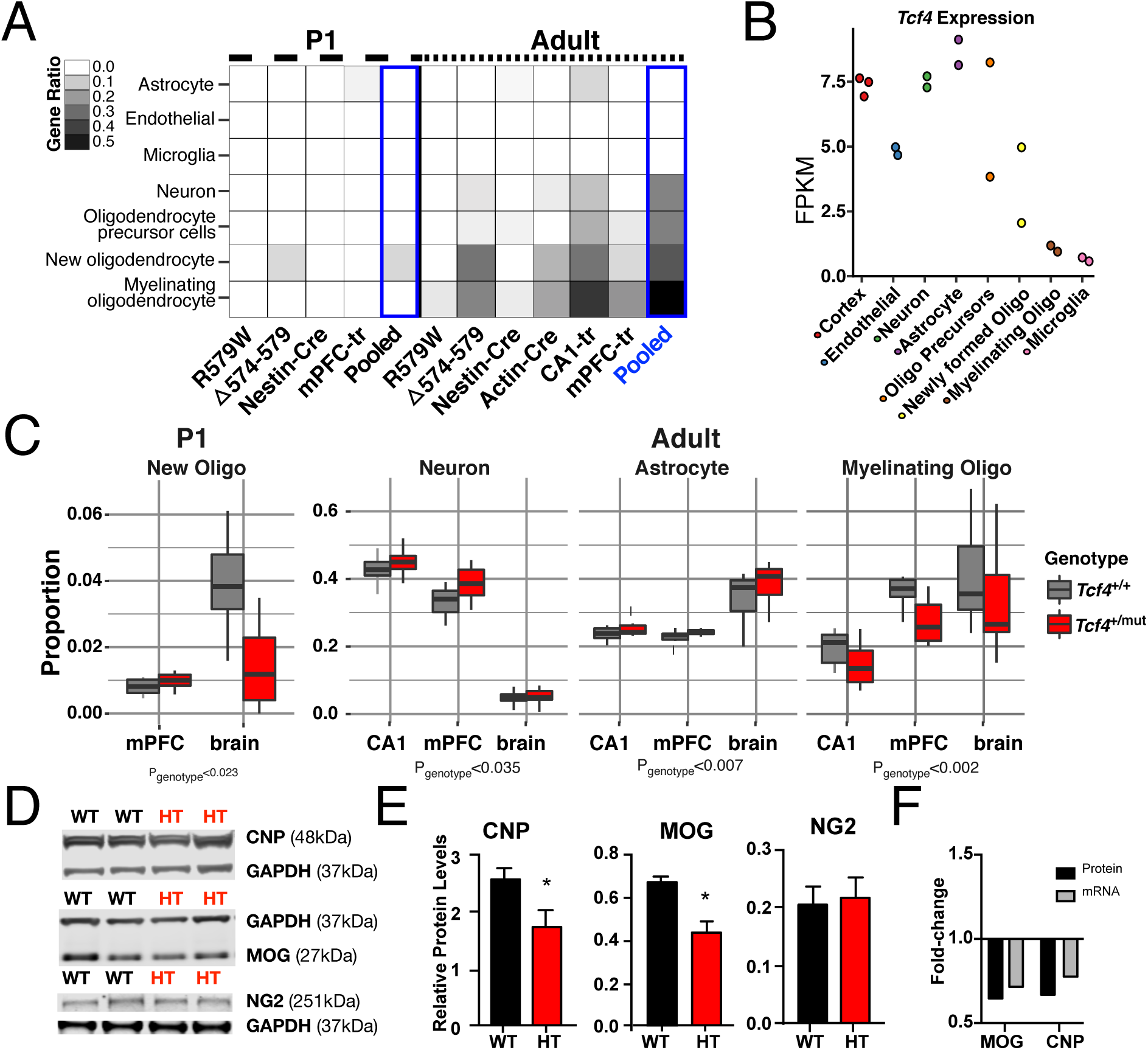
Oligodendrocyte-specific deficits in PTHS model mice. We used RNA-seq data from purified mouse brain cell-types(*14*) with the CIBERSORT(*24*) to predict cell type-specific differences in the *Tcf4*^+/mut^ (PTHS) mice. **(A)** Heatmap plotting the ratio of cell type-specific genes that are DEGs. Differential expression in all adult mouse line were highly specific to myelinating oligodendrocyte signature genes (P_adj_<0.05). New oligodendrocytes, their precursors, neurons, and astrocytes are enriched in DEGs of different forms of *Tcf4* mutations, but most present in the adult brain (P_adj_<0.05). **(B)** *Tcf4* expression in these cell types from Zhang *et al.* **(C)** CIBERSORT cell proportions analysis of *PTHS* mice stratified by sample tissue source. New oligodendrocyte proportions are down in P1 brains, neuron and astrocyte proportions up in adult brain, and myelinating oligodendrocyte proportions are down in adult brain (p<0.05). **(D)** Western blot for myelinating oligodendrocyte proteins, CNP and MOG, and a oligodendrocyte precursor cell protein, NG2, normalized to GAPDH in mPFC-*Tcf4*^+/tr^. **(E)** Relative protein levels of MOG and CNP are significantly decreased in *Tcf4*^+/tr^ brain (unpaired t-test, N = 12 each, p_MOG_=0.0008, p_CNp=_0.0436). Relative levels of NG2 are not differential between genotypes (N = 12, p=0.815). **(F)** The decrease fold-change of CNP and MOG proteins validate decreased levels of mRNA from differential expression analysis. (Abbreviations: R579W = *Tcf4*^+/R579W^; Δ574-579 = *Tcf4*^+/Δ574-579^; Nest-Cre = *Nestin-Cre*::*Tcf4*^+/floxed^; Actin-Cre = *Actin-Cre::Tcf4*^+/floxed^; mPFC-tr = *Tcf4*^+/tr^ medial prefrontal cortex; CA1-tr = *Tcf4*^+/tr^ hippocampal CA1 neurons; FPKM = fragments per kilobase per million reads mapped)

**Fig. 3:**
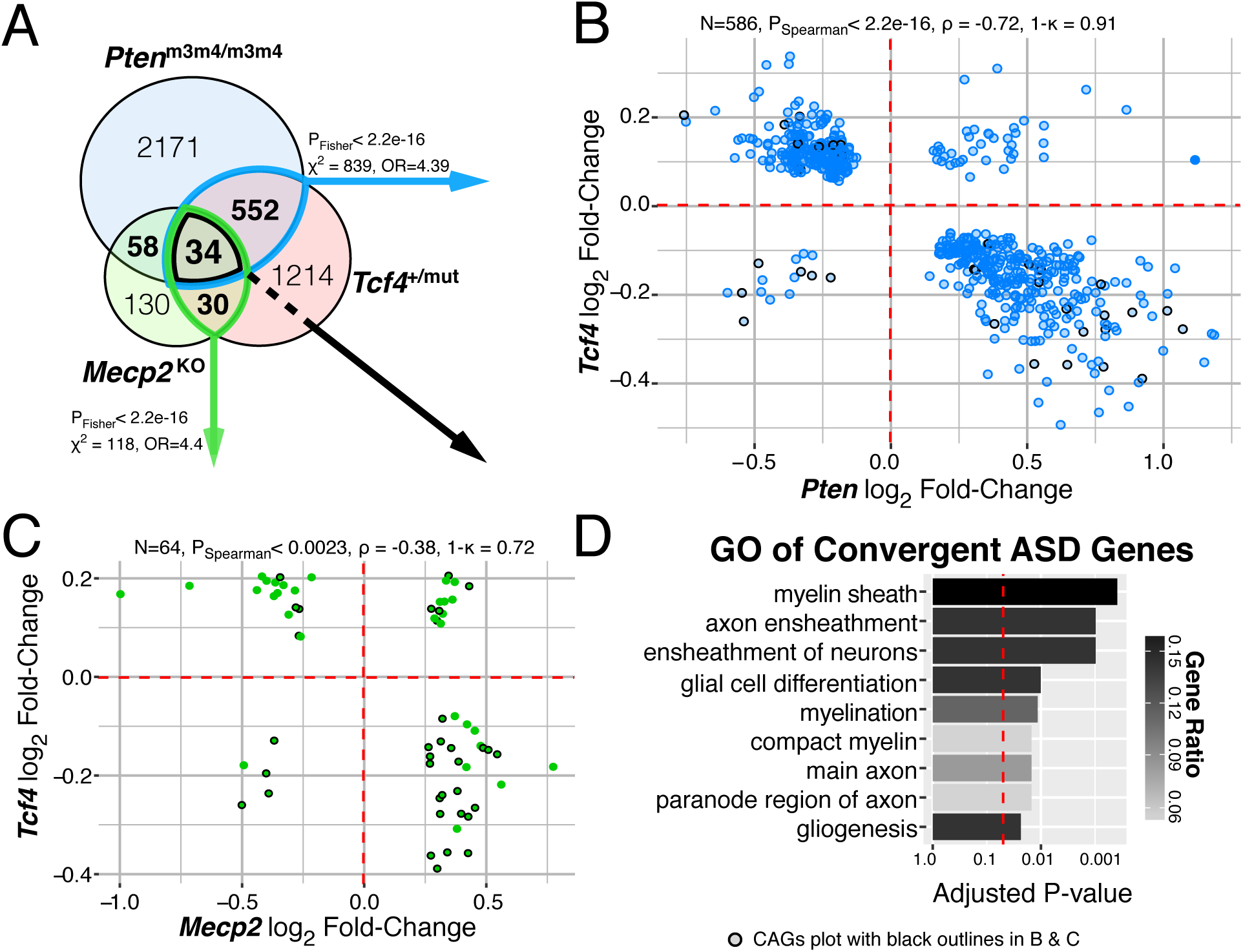
Shared myelination gene regulation between mouse models of syndromic ASD. **(A)** Venn diagram of DEGs in each mouse model of ASD, FDR < 0.05. There is significant overlap of DEGs from *Tcf4* mutation vs. *Mecp2* or *Pten* homozygous mutation (P_Fisher_<2.2e-16). **(B-C)** Log_2_ fold-change comparison of the (N) genes differentially expressed both in *Tcf4* heterozygous mutation and *Mecp2* knockout or *Pten* homozygous mutation. DEG fold-change directionality in TCF4 mutant mice is inversely correlated (ρ < 0) to that in MeCP2 and Pten mutation suggesting TCF4 plays an opposite role in regulating these genes. 34 genes differentially expressed in all three mutations, referred to as the convergent ASD genes (CAG) are plot with black outlines. **(B)** The 586 DEGs in the *Tcf4* vs. *Pten* comparison had 91% opposite fold-change directions and displayed strong negative correlation (P_Spearman_<2.2e-16, ρ = -0.71). **(C)** The 64 overlapping DEGs in the *Tcf4* vs. *Mecp2* group had 72% opposite fold-change directions with significant negative correlation (P_Spearman_<0.0023, ρ = -0.38). **(D)** Top GO terms of the CAGs enrich for myelination processes (P_adj_<0.05). (ρ, Spearman’s correlation coefficient, ρ, concordance rate).

Syndromic forms of ASD are caused by varying mutations yet share significant overlap in their symptomology, suggesting that common biological processes underlie many forms of ASD. We therefore compared DEGs in adult *Tcf4*^+/mut^ mouse brains to mouse models of PTEN-associated autism (*Pten*^m3m4/m3m4^; (*15*, *16*)) and Rett syndrome (*Mecp2^KO^*; (*17*), (*18*)) to characterize shared gene regulation between these three mouse models of syndromic ASD. Remarkably, we found significant overlap of DEGs in *Tcf4*^+/mut^ vs. *Pten*^m3m4/m3m4^and *Tcf4*^+/mut^ vs. *Mecp2^KO^* mutations (Fig. 3A, p < 2e-16), with *Tcf4* mutations inversely regulating genes differentially expressed in *Pten* and *Mecp2* mutations. Overall, 34 DEGs (each at FDR<0.05) were common to all three mouse models, namely the convergent ASD genes (CAGs, Fig. 3A). The CAGs were strongly enriched for several myelination-related GO terms, showing up to 15% of the genes involved in some processes (Fig. 3D, p_adj_<0.0149). The CAGs are generally down-regulated in the *Tcf4* mice while being up-regulated in *Pten* and *Mecp2* mutations, which is consistent with prior studies demonstrating decreased PTEN and MeCP2 protein expression promote oligodendrocyte expansion and maturation(*19*, *20*) (black-outlined points in Fig. 3B-C). Interestingly we observed that *Mecp2* expression was significantly upregulated in the adult PTHS brain (FDR=0.002), and may potentially provide a mechanistic explanation for the observed inverse fold-changes between *Tcf4* and *Mecp2* mutations and their subsequent effects on myelination. *Pten*, however, was not differentially expressed in PTHS mice, so the strong inverse correlation between *Tcf4* and *Pten* mutations may arise from regulation of common downstream targets such as *Akt2* or *Mtor*, two DEGs in our dataset that participate in the Akt-PTEN-mTOR pathway(*21*). The significant overlap of differential expression between these mouse models of syndromic ASD and their apparent correlational effects on myelination suggests deficits in oligodendrocyte development and function is potentially a common molecular pathway disrupted in syndromic ASD.

We examined the potential role of oligodendrocyte disruption in idiopathic ASD, which are more prevalent in the human population, but due to their polygenic nature are more difficult to model in animals. First, we found significant enrichment of adult *Tcf4*^+/mut^ mice DEGs in the SFARI database of scored ASD candidate risk genes. One quarter of the homologous SFARI genes were DEGs in the *Tcf4*^+/mut^ mice (183/583) and 25 of those cause syndromic forms of ASD (Table S5, p_Fisher_=6.36e-14), suggesting TCF4 may be an upstream regulator of a variety of previously identified ASD risk genes. Next, we identified significant enrichment of DEGs in the *Tcf4*^+/mut^ mice among published WGCNA co-expression modules from microarray of human ASD postmortem brain(*22*), BrainSpan RNA-seq of developing human neocortex(*11*), and RNA-seq from human postmortem prefrontal and temporal cortex(*23*) (Fig. 4C). Specifically, we found the strongest enrichment of *Tcf4*^+/mut^ DEGs among human genes involved in myelination, axon ensheathment, and gliogenesis (asdM14, adjusted p=5.6e-22). In addition, we observed *Tcf4*^+/mut^ DEGs showed enrichment in other previously identified co-expression modules involved in synaptic transmission and mRNA processing (Fig. 4C). Enrichment in these gene co-expression modules supports the idea that gene networks disrupted in *Tcf4*^+/mut^ are similar to gene networks disrupted in human ASD, and further identifies myelination as an additional pathway disrupted in ASD.

**Fig. 4:**
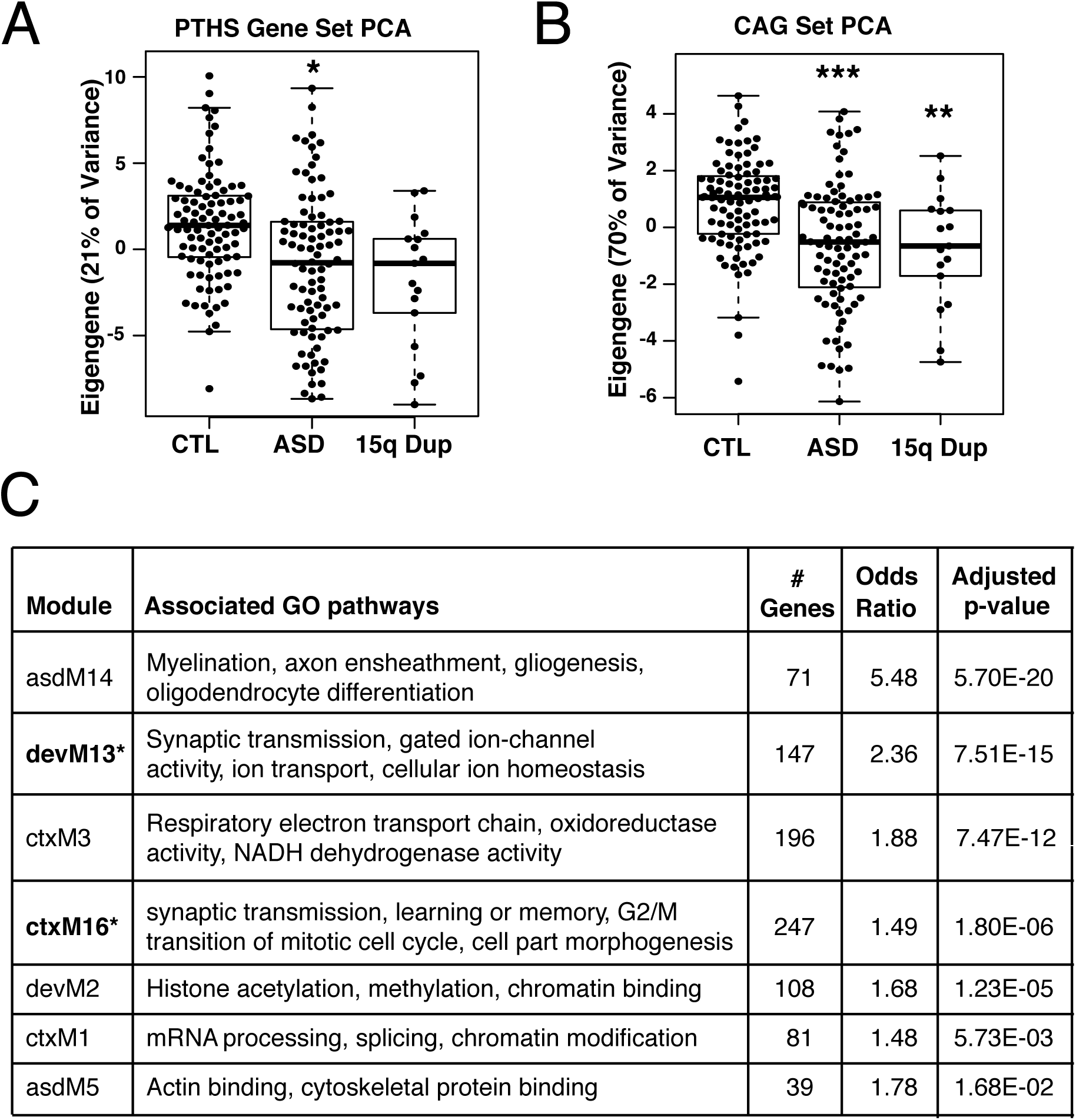
Human-mouse convergence of gene expression in idiopathic and syndromic ASD. **(A)** The eigengene from principle component analysis (PCA) of 1718 human homologs of adult PTHS mouse DEGs in idiopathic ASD and 15q duplication postmortem brain RNA-seq. The PTHS eigengene explains 21% of the gene expression variance and is slightly associated with the ASD diagnosis (p=0.022). **(B)** The eigengene of the CAGs found across the three models of syndromic ASD. The CAG eigengene explains 69% of the gene expression variance and is highly associated both ASD and 15q duplication diagnoses (p_ASD_=1.5e-6, p_15qDup_=0.0098). **(C)** DEGs in adult *PTHS* mice enrich for co-expression modules identified in microarray of human ASD(*22*) (asdM), neocortex development(*11*) (devM), and RNA-seq of human ASD and 15q duplication in cortex(*23*)(ctxM). Table shows the number of DEGs participating in each module, the enrichment odds ratio, and FDR-adjusted p-value.* denotes gene co-expression models previously identified to be associated with ASD patient diagnosis or risk genes.

To better characterize phenotypes identified in mouse models with human illness, we directly compared RNA-seq data between *Tcf4*^+/mut^ mice and postmortem human brain tissue from ASD and 15q duplication postmortem brain from frontal cortex, temporal cortex, and cerebellar vermis(*23*) (Fig. S4A). Differential expression between PTHS mouse and human ASD or 15q duplication was significantly concordant (Fig. S4B, p_adj_<0.05). *Tcf4*^+/mut^ DEGs strongly overlapped with human ASD and 15q duplication at replication p<0.05, displaying slightly more overlap with the 15q duplication diagnosis (Fig. S4C, p_adj_<0.05). Comparing replication of DEGs with a more stringent cutoff (p<0.01), gene regulation became more correlated with higher concordance rates, suggesting these groups of genes better represent the shared pathways affected in both PTHS and human ASD. GO analysis of DEGs in frontal and temporal cortex overlapping with PTHS mouse DEGs enriched for processes related to axon and dendrite projection development and postsynaptic regulation and signaling (Fig. S4E). Lastly, we evaluated the ability of our mouse gene sets to distinguish patients with ASD and 15q duplication from unaffected individuals. We obtained only marginal separation of patients with ASD from controls among the 1718 human homologs of *Tcf4*^+/mut^ mouse DEGs (via the top eigengene, Fig. 4A, p=0.022). However, we significantly separated both patients with ASD and 15q duplication from controls using the CAG set at similar magnitudes (via the top eigengene, Fig. 4C, p_ASD_=1.5e-6, p_15qDup_=0.0098). This top eigengene was directionally consistent between the human and mouse samples, such that decreasing expression of the convergent mouse gene set corresponded to decreased expression of oligodendrocyte genes in human patients with ASD.

By comparing differential expression in mouse models of *Tcf4*^+/mut^ mice with human ASD and 15q duplication, we observed similar trends in differential expression between the datasets, particularly among the CAGs. The observed positive correlation between altered gene regulation in mouse models of syndromic ASD with those in idiopathic human ASD strongly supports the hypothesis that defects in myelination are a common pathophysiology in syndromic and idiopathic ASD. This cross-species framework demonstrates the usefulness of mouse models to human ASD and highlights novel targets and pathways for potential pharmaceutical intervention.

## Acknowledgements

We are grateful for the vision and generosity of the Lieber and Maltz families, who made this work possible. This work was supported by the Lieber Institute, the Pitt-Hopkins Research Foundation Award (B.J.M, B.D.P., C.T., D.S., and A.J.K), an NIMH R56MH104593 grant (B.J.M.), an NIH R01 grant 5R01MH110487 (B.J.M), a Johns Hopkins PURA grant (B.N.P), a UPenn Orphan Disease Center Million Dollar Bike Ride grant MDBR-15-108-PH (B.D.P. and C.T.), an NARSAD Young investigator grant #20653 from the Brain Behavior Research Foundation (C.T.), an NINDS grant P30NS045892 (J.M.S.), an NICHD grant P300HD03110 (J.M.S), and an NIH R01 grant MH104158 (D.S. and A.J.K). The authors declare that they have no competing financial interests.

## Supplemental Materials

### Materials and Methods

#### Animals and tissue collection

The *Tcf4*^+/tr^ mouse model of PTHS(*6*) were heterozygous for a deletion of the DNA-binding domain of the Transcription Factor 4(B6;129-TCF4tm1Zhu/J; stock number 013598, Jackson Laboratory, Bar Harbor, ME)(*8*). This deletion truncates the transcript after the first helix of the basic helix-loop-helix (bHLH) domain in exon 19(*8*). This mouse colony was backcrossed 6 generations, maintained by SoBran on a 12-hour light cycle, and fed *ad libitum*. *Tcf4*^+/tr^ mouse samples were matched with WT littermates, and sex was randomly selected in each genotype and age group. The *Tcf4^R579W^* and *Tcf4^Δ574-579^* were generated using Crispr/Cas9 technology by the Animals Models Core facility at UNC. Heterozygous founders from each mutation were checked for off-target effects of the guide RNA, to which no changes were found. The *Tcf4^+/floxed^* mice were generated previously (Berqvist et al., 2000), and the *Actin-Cre* (JAX stock # 019099) and Nestin-Cre (JAX stock #003771) mice were purchased from Jackson Laboratories. The *Tcf4^+/R579W^*, *Tcf4^+/Δ574-579^*, *Actin-Cre::Tcf4^+/floxed^*, and *Nestin-Cre::Tcf4^+/floxed^* mice were maintained on a congenic C57/BL6 background, and maintained on a 12:12 light dark cycle with *ad libitum* access to food and water. Age-matched, sex-matched WT littermates were used as controls for all samples.

#### qRT-PCR

To measure *Tcf4* transcript expression across *Tcf4*^+/tr^ mouse lifespan in mouse, three cortical samples were collected each from both genotypes over 11 developmental ages from embryonic day 12 (E12) to postnatal day 42 (P42, adult). Embryonic samples required micro-dissection for frontal cortical tissue. All samples were separately flash frozen and homogenized in Trizol (Life Technologies). Aqueous phase was mixed with a 1:1 volume of 70% ethanol prior to purification using RNeasy mini columns treated with DNAse according to manufacturer’s protocol (Qiagen). RNA samples were then reverse-transcribed to cDNA using the Quantscript Reverse Transcriptase kit (Qiagen). Amplification of cDNA was performed with iTaq SYBR Green Supermix (Bio-Rad) and *Tcf4* primers were designed using with Primer 3 software (http://bioinfo.ut.ee/primer3-0.4.0/). Primers were designed to span exons 19 and 20 to measure the full-length transcript. End-point PCR followed by product sequencing, in addition to cDNA dilution series and melt curve analysis, were used to verify primer design efficiency and specificity. Real time PCR was performed on the 7900HT Fast Real-Time PCR system (Applied Biosystems) and melt curve analysis was done for absolute quantification of primer efficiency. Data was expressed as fold-change of gene of interest normalized to *Gapdh* expression using 2^-delta delta Ct. Lifespan expression fold-change was analyzed between the two genotypes with a two-way ANOVA with Sidak’s multiple comparison test for post hoc analysis in the GraphPad’s Prism software.

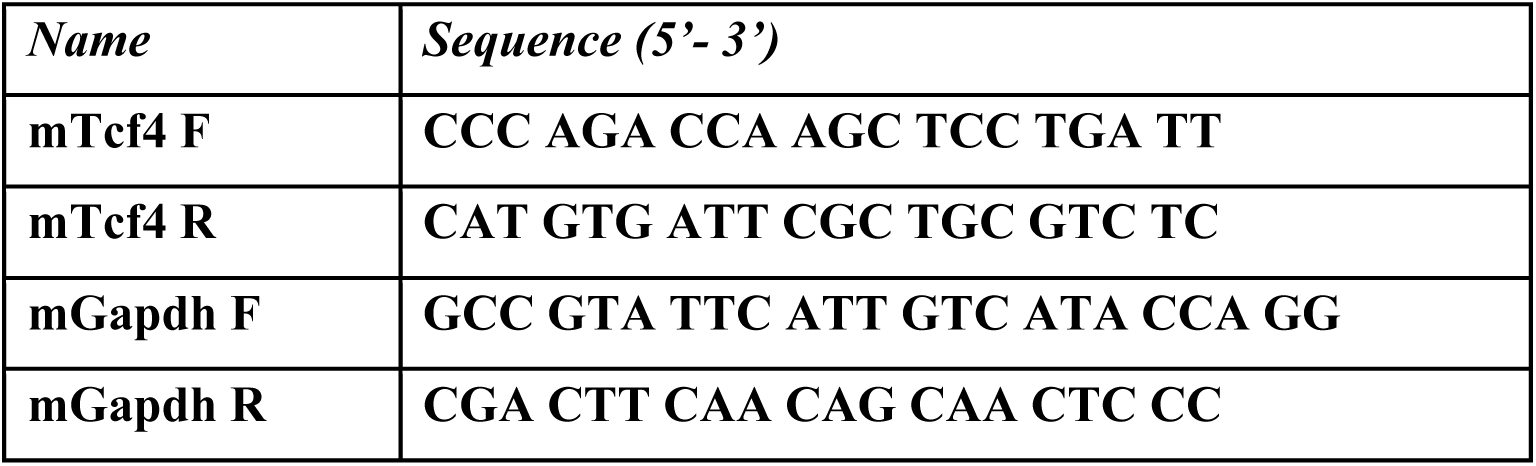

#### Western Blot

TCF4 protein levels were measured across development in E12, P1, and adult ages (N=3 per condition). Mice cortices were extracted and flash frozen on dry ice. Dissected tissue was homogenized with T10 basic ultra-turrax (IKA) in a 1:15 ratio of tissue (mg): RIPA buffer containing protease inhibitor cocktail (Amresco). Samples were sonicated for 20 cycles using Sonifier 250 (Branson Ultrasonics) set at an output control of 1.5, and duty cycle of 60.05. Lysates were mixed with 20% SDS (Amresco) for a final concentration of 2% and resonicated. Lysates were then incubated on a rotator for 1hr in 4°C and centrifuged at 20,000g for 5 minutes at 4°C. Supernatants were quantified using BCA kit (Pierce).

Samples were blinded and randomly ordered on Western blots to control for batch bias. Total protein amounts of 20μg were used from each sample and separated using a 4-12% gradient Novex Bis-Tris Bolt SDS-PAGE gel via gel electrophoresis and transferred to 0.45 μm nitrocellulose membranes. Membranes were incubated at room temperature for 1hr in Odyssey PBS blocking buffer (Li-Cor), probed with anti-ITF-2 (N-16) (1:500, Santa Cruz) and anti-GAPDH (1:1000, Abcam) primary antibodies in Odyssey PBS blocking buffer overnight at 4°C, and detected using IRdye donkey anti-goat 680 (1:10,000, Li-Cor) and IRdye donkey anti-rabbit 800 (1:10,000, Li-Cor). Antibody detection and quantification was carried out using the LI-COR Odyssey infrared system and software. The *Tcf4*^+/tr^ mouse produces two bands for the full-length (∼75kDa) and truncated protein (∼70kDa) variants. We compared protein levels of the full-length protein over time and between genotypes using a two-way ANOVA with Sidak’s multiple comparison test for post hoc analysis in Prism.

CNP, MOG, and NG2 protein levels were similarly extracted and measured from whole brain lysates of *Tcf4*^+/tr^ and WT adult mice (N=6 per genotype). Starting amounts of protein lysates (30μg, 20 μg, and 60ug) were respectively probed with 1:1000 dilutions of each primary antibody (mouse anti-CNP, Millipore; mouse anti-MOG, Abcam, and rabbit anti-NG2, Millipore) and GAPDH and detected with 1:10,000 dilutions of Li-Cor IRdye (donkey anti-mouse 680, donkey anti-rabbit 800). Levels of CNP, MOG, and NG2 were normalized to GAPDH. We compared relative protein levels between genotypes using the student’s t-test in Prism.

#### Library preparation and RNA sequencing

RNA was extracted from the brains of *Tcf4*^+/tr^ and WT littermates at two developmental time points, postnatal day 1 and >42 with 6 animals per group (total N=24). Animals were collected at the indicated time point, euthanized, and brains removed from the skull. The medial prefrontal cortex (mPFC) was rapidly dissected on an ice block, and the tissue samples were immediately subjected to RNA purification. Sequencing libraries were prepared from RNA from each mouse cortex using the Illumina TruSeq Stranded RNA HT sample preparation kit with Ribo-Zero^TM^Gold. Libraries were barcoded and then sequenced using an Illumina HiSeq 3000 at 100bp paired-end reads for targeted coverage of 50 million sequencing fragments (100 million reads) per sample in the LIBD Sequencing Core Facility. There was a median of 129M(Million) mapped reads per sample (interquartile range, IQR: 109M-140M), of which a median 52% (IQR: 49%-54%) were assigned to genes (based on exonic sequence overlap).

Brains collected from *Tcf4^+/R579W^*, *Tcf4^+/Δ574-579^*, *Nestin-Cre:: Tcf4^+/Flox^*, *Actin-Cre::Tcf4^+/Flox^*, and littermate controls were dissected at either P0-P2 or P60-P80. Brains were rapidly dissected, flash frozen in a dry ice and ethanol bath, and stored at -80°C. RNA was extracted from one (P60-P80 mice) or both (P0-P2) cerebral hemispheres using the RNeasy Plus kit (Qiagen) per manufacturer’s instructions with the following modifications. The frozen tissue was thawed on ice and then hand homogenized in 500μl of Buffer RLT^+^ on ice. The crude RNA lysates were diluted with additional Buffer RLT^+^ to a final volume of 2mL. For purification of RNA, 400μl of P60-P80 lysate or 200μl of P0-P2 lysate were used from the total RNA lysates (2mL). The crude RNA lysate was twice passed through the genomic DNA eliminator column, and the purification proceeded according to the manufacturer’s instructions (Qiagen). A 40μl aliquot of purified RNA was used for additional purification using the RNA Clean and Concentrator kit (Zymo). Briefly, the purified RNA was incubated with RNase-free DNase I (Zymo) for 15 minutes at room temperature. The digested RNA was then washed and cleaned on column per the manufacturer’s instructions. All tested samples had 260/280 and 260/230 ratios ≥ 2.0 measured using a NanoDrop (Thermo Scientific). All RNA samples were verified to have an RNA integrity number (RIN) INNA integrity number on the TapeStation 2000 (Agilent Technologies). Unstranded library construction, quality control, and RNA-sequencing were performed by Beijing Genomic Institute (BGI, Beijing, China) using the Illumina HiSeq4000 at single-end 50bp reads. These samples had a median of 29M reads mapped per sample (IQR: 26M-31M), of which a median 81% (IQR: 79%-83%) were assigned to genes (based on exonic sequence overlap).

#### Read mapping and quantification of gene expression

HISAT2 genome indices for mm10/GRCm38.p4 were created according to developer’s instructions. Reads were aligned to the mm10 mouse genome using the splice-read mapper HISAT2 (version 2.0.4)(*25*) using the reference transcriptome to initially guide alignment, based on known transcripts of the GENCODE build [hisat2 -p 8 -x $GRCm38index -1 $FP -2 $RP –S $SAM --rna-strandness RF --phred33 2> $SUM]. Single-end reads were aligned with the argument [-U $UP]. Unstranded reads did not include the [--rna-strandness] argument. Gene expression levels were calculated with the featureCounts tool (version 1.5.0) based on the GENCODE version M11 annotations of the mm10 genome. *Tcf4*^+/tr^ expression was summarized with featureCounts arguments for reversely-stranded read pairs [featureCounts -s 2 -p -T 8 –a $GTF -o $OUT $BAM]. Other *Tcf4*^+/mut^ samples were prepared with unstranded library kits, so expression was summarized with [-s 0] argument instead. ($FP: forward pair; $RP: reverse pair; $SAM: output alignment file, RF: HISAT argument for reversely stranded library prep; $UP: unpaired reads)

#### Differential expression across *TCF4* mutations and brain regions

Differential expression in P1 and adult mice brain from *Tcf4* mutation was determined by pooling samples from medial PFC, whole brain, and hippocampal CA1 from 5 forms of *Tcf4*^+/mut^ (N_P1_=28, N_Adult_=69). The R package DESeq2(*26*) used raw gene counts to determine DEGs by genotype with the linear model [geneCounts ∼ Genotype + Line + geneAssignmentRate+ SVs]. Lowly expressed genes were dropped from differential expression (average normalized counts across all samples less than 1). Expression was adjusted for mouse line and gene assignment rates to account for differences between mouse lines, animal care, tissue source, library preparation, and sequencing procedures. Surrogate variables (SV) determined by the R package sva(*27*) were included to remove batch effects and noise in gene expression data from unknown or un-modeled sources of variation. One low quality adult *Tcf4*^+/tr^ sample was identified in hierarchal clustering and was dropped from analysis. The p-values were adjusted for multiple testing through DESeq2 with a target alpha = 0.05, and mouse genes were considered DEGs at FDR<0.05.

#### Comparison between differential expression analyses

Sub-analyses to determine DEGs within each mouse line and age group were similarly determined with DESeq2 with the linear model [geneCounts ∼ Genotype + SVs] at alpha = 0.05. Samples within these groups are considered balanced and matched. A DEG from one dataset is considered ‘replicated’ if it is differentially expressed at p<0.05 in more than one dataset (Fig. 4, 5, and S1).

#### Functional gene set analysis on the DEGs

We found enriched gene pathways in GO databases with the R-package clusterProfiler(*28*). The clusterProfiler analysis tested the DEGs at p<0.01 from the pooled analysis in P1 and adult samples for over-represented gene sets using hypergeometric tests. DEGs are separated by positive and negative log_2_ fold-change. We defined the background as the list of expressed genes with mean normalized counts > 1 and adjusted for multiple testing with q-value<0.05.

#### Cell type-specific expression and relative proportion analysis

We first used the CSEA approach(*13*) to determine cell type specificity of these *TCF4* mutations, which are based on over-representation of cell type-specific genes, determined by TRAP. Analyses and plots were generated with the online CSEA tool (http://genetics.wustl.edu/jdlab/csea-tool-2/). Each cell type is represented by a 4-level bullseye plot scaled by the number of transcripts unique to that cell type at multiple specificity index thresholds (pSI<0.05, 0.01, 0.001, 0.0001). Enrichment BH-adjusted p-values are plotted in each level of specificity (i.e. the most enriched cell types will have the lowest p-values in all rings). Independent validation of CSEA analysis took RNA-seq raw gene counts from purified cell types of mouse brain from Zhang et al.(*14*) and the CIBERSORT online tool (cibersort.stanford.edu) to create the set of signature genes using default parameters. The signature genes for enrichment was further limited to those upregulated in these cell types from DESeq2 analysis (Fig. 3A). Raw gene counts from *PTHS* mice were passed to CIBERSORT to predict relative proportions of each cell type. Change in proportions of each cell type was determined by fitting the linear model ‘Proportion∼ Genotype +Line’.

#### Comparing transcriptomes of multiple syndromic ASD mouse models

We compared DEGs from *Tcf4*^+/mut^ mice with two mouse models of ASD, *MeCP2* knockout (*Mecp2*^KO^)(*18*) and homozygous *Pten* mutation (*Pten*^m3m4/m3/m4^)(*16*). We processed the *Mecp2*^KO^ and *Pten*^m3m4/m3/m4^ RNA-seq datasets as described below and compared genes differentially expressed in our pooled TCF4 analysis to those also differentially expressed in MeCP2 and Pten homozygous mutations (FDR<0.05). We tested enrichment of differential expression, log_2_ fold-change correlation, and log_2_ fold-change directionality concordance with Fisher’s Exact test and Spearman correlation test. We also report ρ, the Spearman’s correlation coefficient, and 1-κ, the rate of genes with opposite log_2_ fold-change directionality. GO analysis for Fig. 3D was performed on the set of genes differentially expressed in all three mouse models.

#### Comparing to human ASD risk genes, expression networks, and differential expression

To directly compare our PTHS mice models with sporadic human ASD, we compared our DEGs to the Simons Foundation Autism Research Initiative (SFARI) ASD risk gene database(*29*) and we searched for overlap using weighted gene co-expression network analyses (WGCNAs) from human ASD and neocortical development, as well as gene expression of a recent large RNA-seq study on human ASD postmortem brain(*23*). We downloaded the list of ASD risk genes scored by SFARI by the evidence of their risk associations to idiopathic or syndromic ASD. We included 583 of the 632 risk genes that were homologous mice genes that are expressed in our RNA-seq dataset and used the Fisher’s Exact test to test for over-representation of our DEGs in the list of human ASD risk genes. We tested enrichment with weighted gene co-expression network analysis (WGCNA) of human brain development(*11*), microarray of human ASD(*22*), and RNA-seq of human ASD(*23*). For WGCNA data from the Voineagu *et al.* study, we assigned genes to the module that they have strongest membership (kME > 0.7). Here, we limited analysis to only expressed genes in the mouse data that have human homologs present in each WGCNA study. As described in these papers, we tested enrichment in each co-expression module with two-tailed Fisher’s Exact test, included only positive enrichment (odds ratio > 1), and adjusted for multiple testing with the false discovery rate (FDR<0.05). We also provide a companion GO analysis of these enriched modules to interpret the molecular pathways involved in each co-expression networks. We used all genes in each module with the background as the list of genes reported in each study.

#### Comparing mouse models of PTHS with human ASD RNA-seq

Lastly, we compared adult PTHS DEGs with differential expression of human ASD vs. control in three brain regions (DLPFC, auditory cortex, and cerebellar vermis). We used biomaRt(*30*) to map mice genes to their human homolog genes and determined fold-change concordance with chi square test of independence in significant DEGs in both datasets (FDR<0.01 in mice, p<0.05 in human). We tested the log_2_ fold-change correlation with Fisher’s Exact tests. We report κ, the rate of genes with concordant log_2_ fold-change directionalities between PTHS mouse and human ASD or 15q duplication in each brain region with their respective permutation p-values of 1000 iterations. We subsequently performed GO analysis on genes in both mouse and human for each brain region (as above) with the background as the set of expressed genes with mouse homologs.

#### Other RNA-seq data processing

Kennedy *et al.* RNA-seq data from hippocampal CA1 of *Tcf4*^+/tr^ mice(*12*): We acquired from the authors 16 BAM files from RNA-seq of CA1 pyramidal neurons of unconditioned Tcf4^+/tr^ and WT P60 mice. Reads were extracted from the BAM files to be realigned as single-end unstranded reads with HISAT2. Expression was summarized with featureCounts calls for single-end unstranded reads. Differential expression by genotype adjusting for surrogate variables was determined with DESeq2.

Gabel *et al.* RNA-seq data of MeCP2 homozygous knockout mice(*18*): We downloaded the MeCP2 dataset from accession GSE60077 (N=6, WT vs. homozygous knock-out, visual cortex). Single-end unstranded raw RNA-seq reads were aligned to the genome with arguments for single-end unstranded. FeatureCounts with similar arguments were used to quantify expression of genes [featureCounts -a $GTF -o $OUT $BAM]. Differential expression was determined in knockout vs. WT mice adjusting for surrogate variables.

Tilot *et al.* RNA-seq data of Pten homozygous mutation mice(*16*): We downloaded the Pten dataset from accession GSE59318 (N= 6, WT vs. homozygous mutants, P42 weeks old). Reversely stranded, paired-end RNA-seq reads were aligned to the genome. featureCounts was used to quantify expression of genes. Differential expression of homozygous mutant mice vs. WT was determined using surrogate variables.

Zhang et al. mouse brain cell type RNA-seq data(*14*): We downloaded the Zhang dataset from accession GSE52564 (N=17, 7 purified cell types and 1 bulk tissue). Reversely stranded, paired-end RNA-seq reads were aligned to the genome. FeatureCounts was used to quantify expression of genes.

Parikshak et al. human ASD RNA-seq data(*23*): We acquired RNA-seq reads of postmortem human brain from the authors, and aligned to the human genome hg38 and GRCh38 gene annotations with HISAT calls for paired, reversely stranded reads. Reads were summarized with featureCounts with analogous parameters. We further summarized library normalized sum coverage with bwtool to measure expression of degradation prone expressed regions from RNA-seq libraries prepared with Ribozero for the qSVA method explained in Jaffe et al, (in review). This degradation region expressed matrix is used in the qSVA analysis method to account for latent degradation effects not completely modeled with RIN. Differential expression in each brain region was then determined with DESeq2 with the statistical model ‘Expression∼ Diagnosis + geneAssignmentRate + SequencingBatch + BrainBank + RIN + Age + Sex+ qSVs’.

#### RNA-seq data accession and code availability

RNA-seq data from cortex and whole-brain of TCF4 mutant mice are available at {Accession codes}. Intermediary files or analyses generated with our RNA-seq processing pipeline are available upon request from authors. R code used to analyze data in this study and analyzed data are available at github.com/LieberInstitute/PTHS_mouse.

#### Author Contributions

B.N.P. and A.E.J. performed RNA-seq analysis; S.C.P. collected samples and contributed to experimental design; M.N.C. performed RT-qPCR and Western Blot experiments; J.F.B. and B.C.M. performed Western Blot and IHC experiments; C.T., J.M.S., A.J.K, D.S., and B.D.P contributed RNA-seq datasets; J.H.S. performed RNA sequencing; E.E.B contributed to RNA-seq data processing; B.N.P, A.E.J, and B.J.M contributed to experimental design, data analysis, and writing. All authors discussed results and commented on manuscript.

**Fig. S1:**
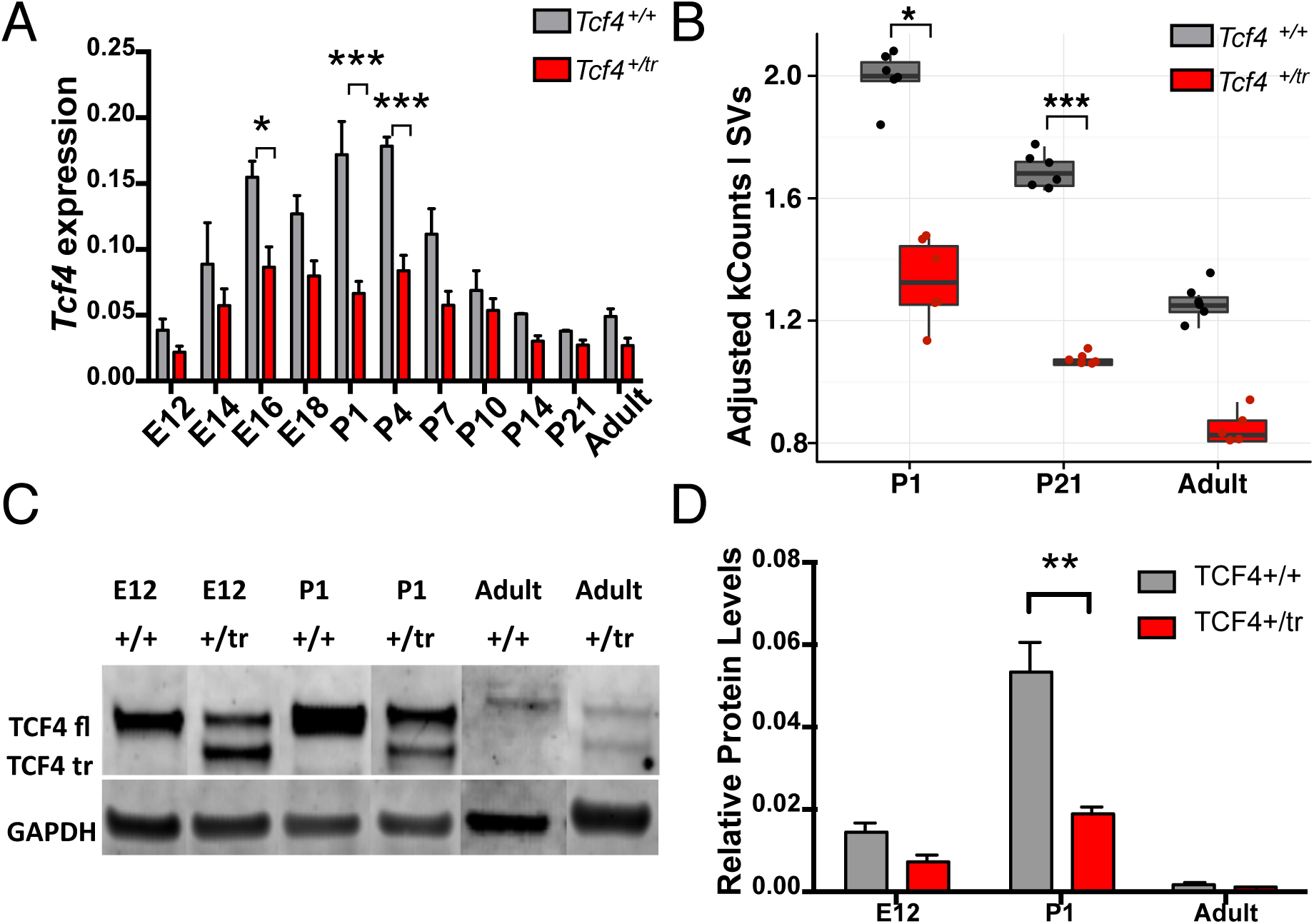
Heterozygous truncation of *Tcf4* decreases levels of *Tcf4* mRNA and protein. (A-B) Comparison of lifespan expression patterns of TCF4 in heterozygous (*Tcf4*^+/tr^) mice and wildtype (*Tcf4*^+/+^) littermates in qRT-PCR and RNA-sequencing analyses. mRNA and protein were extracted from frontal cortex of mice across developmental ages. (A) qRT-PCR analysis of full-length*Tcf4* transcripts from mouse frontal cortex (N=3 per condition) shows *Tcf4*^+/tr^ have the greatest decrease *Tcf4* expression around postnatal days 1-4 (P1-4) (p<0.05). (B) RNA-sequencing analysis also shows TCF4 expression decreased in the *Tcf4*^+/tr^ mouse in the exon after the truncation (N=6 per condition). P1 and P21 *Tcf4*^+/tr^ mice had significant decrease of Tcf4 exons (FDR < 0.05). Shown in (C) is a Western blot of endogenous mouse TCF4 at three ages (E12, P1, and P42). A single full-length (TCF4 fl) protein is observed in lysates from wildtype (+/+) mouse brain. A truncated TCF4 (TCF4 tr) protein is observed in lysates from heterozygous (+/tr) mouse brain. (D) Equal amounts of protein lysate were subjected to Western blot analysis for endogenous TCF4 across development (N=3 per condition). Full-length TCF4 protein is decreased at P1 in the TCF4^+/tr^ mice (p<0.01). (*<0.05, **<0.01, ***< 0.001)

**Fig. S2:**
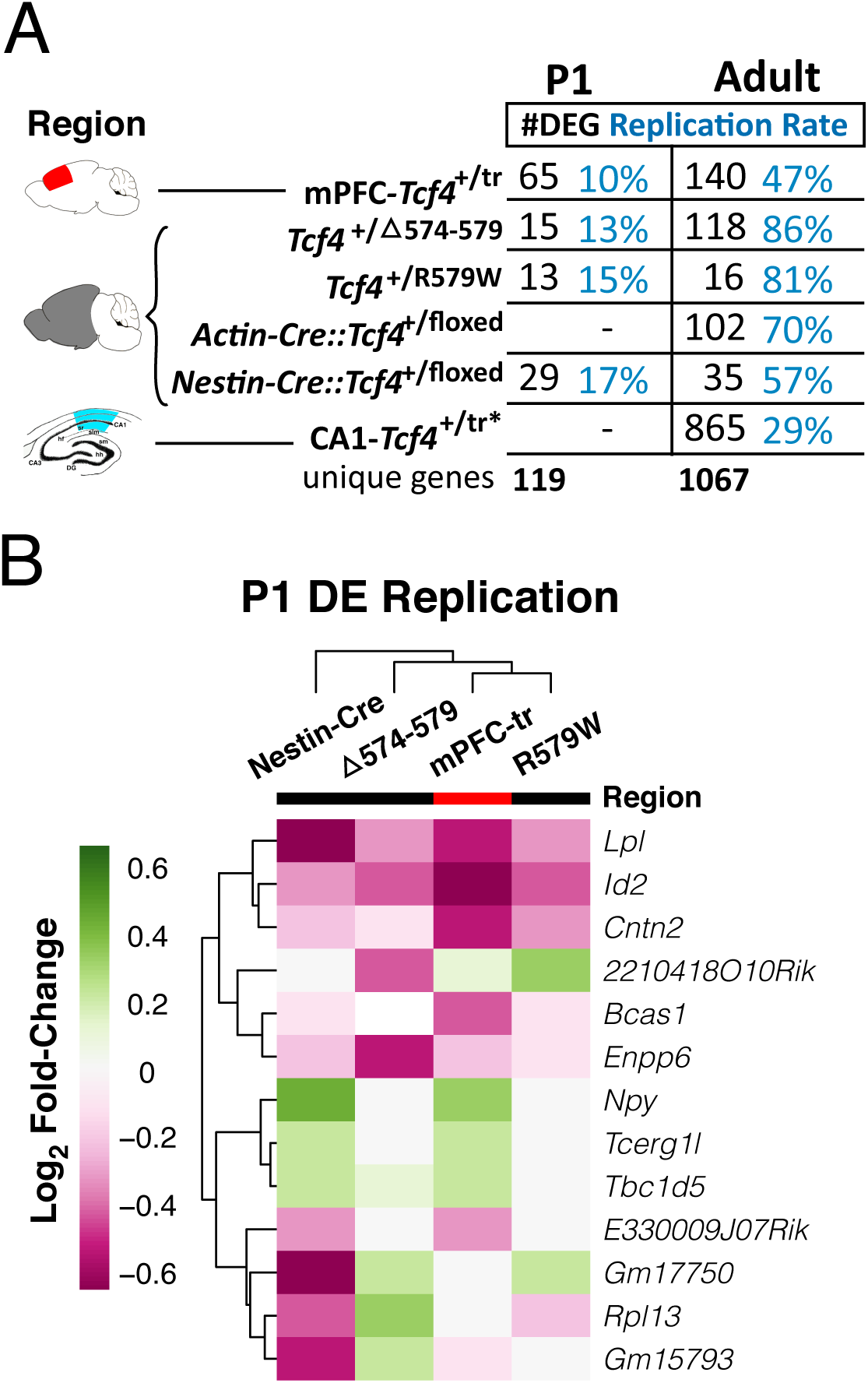
Replicated differential expression across PTHS models. **(A)** Table of DEGs (FDR<0.05) and % of differential expression replication across different forms of *Tcf4* mutation in P1 and adult mice. Most DEGs and replication occur in adult mice. **(B)** Differential expression log_2_ fold-change heatmap comparing replicated DEGs across various models of *Tcf4* mutations in P1 (replication p<0.05).

**Fig. S3:**
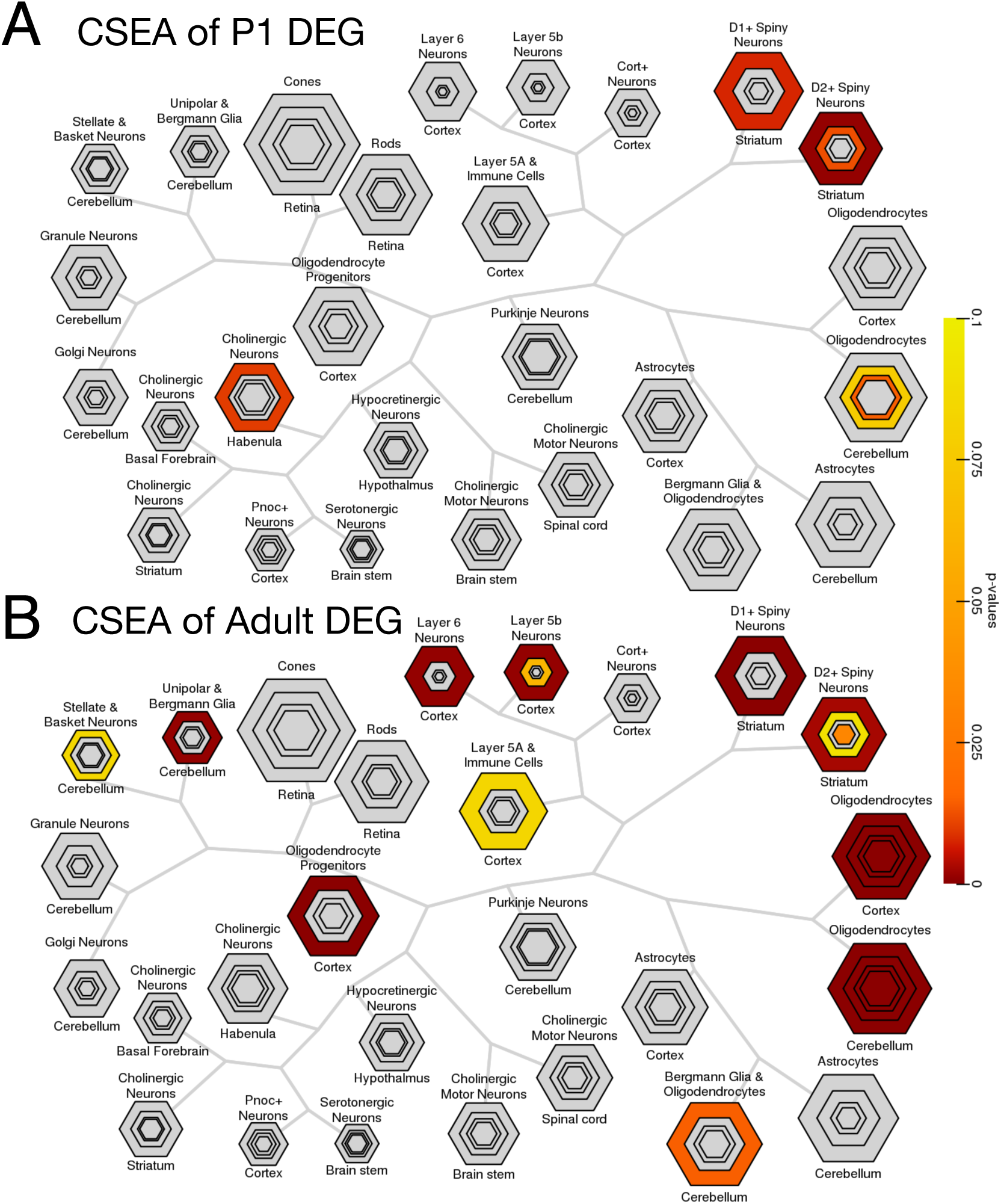
Cell type-specific expression analysis in *PTHS* mice. Bulls-eye plots from CSEA analysis of DEGs in **(A)** P1 and **(B)** adult *PTHS* mice. The bulls-eye plot size is scaled to the number of genes specific to a cell type at increasing levels of specificity as published by Xu *et al., 2014*^13^. The FDR-adjusted p-value is plot for each level of specificity, with unenriched groups colored gray. Cell type bulls-eye plots are arranged by hierarchal distance of their specific gene expression levels. **(A)** P1 DEGs enriched for D1+, D2+, and cholinergic neurons (P_adj_<0.05). **(B)** Adult DEGs strongly enrich for oligodendrocytes among other neuronal cell types.

**Fig. S4:**
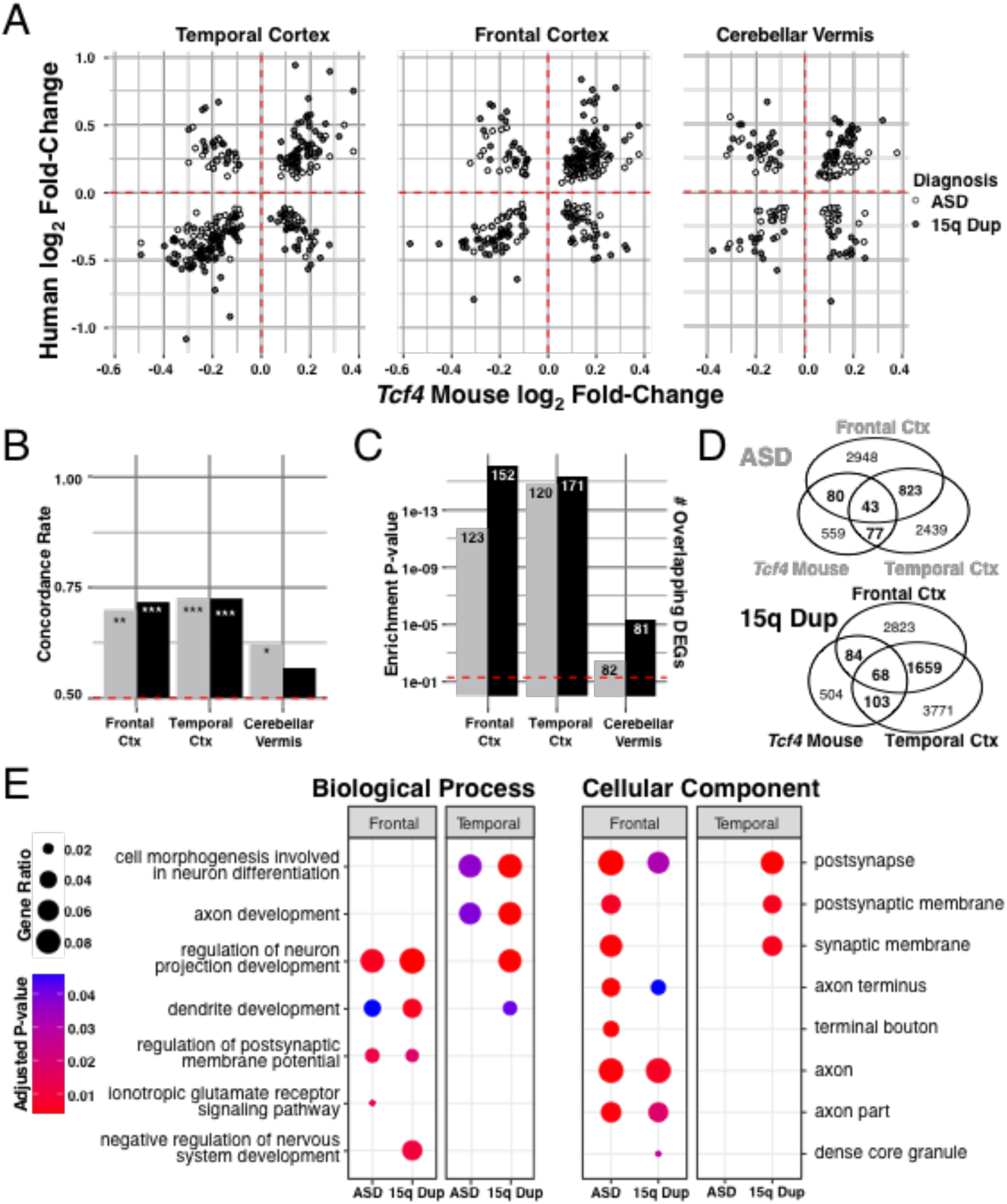
Concordant gene regulation between PTHS mice and human ASD. Comparison of differential expression in adult *PTHS* mice with human ASD and 15q duplication (15q Dup) in postmortem frontal, temporal, and cerebellum. (Human DEGs p<0.05, mouse DEGs, FDR<0.01). **(A)** Log_2_ fold-change comparison of adult PTHS mouse DEGs replicated in human ASD and 15q Dup in each tissue region (p<0.05). Gene regulation in *PTHS* mice cluster closest with ASD differential expression in cortex. **(B)** More than 50% of replicated *PTHS* DEGs had concordant fold-change directionality. Null permutation p-value significance of human-mouse gene fold-change concordance from 1000 permutations are reported (*, p<0.05; **, p<0.01; *** p<0.001). **(C)** Replicated DEGs in ASD and 15q Dup are significantly enriched in all tissues, mostly in the cortex. **(D)** Venn diagram showing overlap of PTHS mouse DEGs with human ASD or 15q Dup in cortical tissues. **(E)** Gene ontology analysis shows tissue-specific biological processes and cellular components between overlap of PTHS mouse and human ASD or 15q Dup. The gene sets are largely brain region specific and concordant between human ASD and 15q Dup.

**Supplemental Table Legends:** (Excel tables attached as a zipped file separately)

**Table S1: Separate differential expression of each *PTHS* mouse line in each age group.** Differential expression in each of the 5 mouse lines of *Tcf4* mutations in postnatal day 1 (P1) or adult mice. Table lists all genes with a least marginal differential expression (p<0.01). Tab names of the table denote the mouse line and age of the samples (Line.Age). Abbreviations: R579W = *Tcf4*^+/R579W^; Del = *Tcf4*^+/Δ574-579^; Nest = *Nestin-Cre*::*Tcf4*^+/floxed^; Act = Actin-Cre:: *Tcf4*^+/floxed^; PFC-tr = *Tcf4*^+/tr^ medial prefrontal cortex; CA1-tr = *Tcf4*^+/tr^ hippocampal CA1 neurons.

**Table S2: Mega differential expression of PTHS mice by age group.** Pooled differential expression analysis by *Tcf4* mutation across the 5 mouse models PTHS by age. Table lists all genes with a least marginal differential expression (p<0.01). Tables include differential expression in P1 and adult mice as well as those genes differentially expressed at both ages.

**Tables S3: Gene ontology analysis in PTHS mice.** GO terms enriched from the pooled differential expression analysis in each age group.

**Table S4: Common DEGs in mouse *Tcf4*, *Pten*, and *Mecp2* mutations.** The 34 genes differentially expressed across the three mouse models of syndromic ASD and enriched gene ontology terms for these genes.

**Table S5: PTHS mouse DEGs in SFARI.** The list of differentially expressed genes in the adult PTHS mice implicated by the Simons Foundation Autism Research Initiative (SFARI) in ASD.

